# PhaseGen: Exact Solutions for Time-Inhomogenous Multivariate Coalescent Distributions under Diverse Demographies

**DOI:** 10.1101/2025.05.13.653888

**Authors:** Janek Sendrowski, Asger Hobolth

**Affiliations:** Bioinformatics Research Center, Aarhus University, 8000 Aarhus, Denmark; Department of Mathematics, Aarhus University, 8000 Aarhus, Denmark

**Keywords:** coalescent theory, phase-type theory, population genetics, demographic inference, software package, time-inhomogeneous coalescent processes, gradient-based optimization, multiple-merger coalescent, site frequency spectrum

## Abstract

Phase-type theory is emerging as a powerful framework for modeling coalescent processes, allowing for the exact computation of quantities of interest. This includes moments of tree height, total branch length, the site-frequency spectrum (SFS), and the full distribution of the time to the most recent common ancestor (TMRCA). However, prior applications have largely been limited to time-homogeneous settings, with constant population sizes and migration rates, restricting the range of demographic scenarios that can be modeled. In this study, we apply time-inhomogeneous phase-type theory to enable the exact computation of (cross-)moments of arbitrary order and reward structure under piecewise-constant demographies. This extension enables the modeling of significantly more complex demographic scenarios, including population expansions, contractions, bottlenecks, and splits. It furthermore supports the fitting of demo-graphic models to data through gradient-based optimization. To support these advancements, we introduce PhaseGen—a software package designed for the numerically stable computation of exact moments under diverse demographic scenarios, with support for gradient-based parameter estimation.

## 1 Introduction

Population genetics modeling software is essential in evolutionary biology, enabling researchers to explore complex demographic scenarios, evaluate evolutionary hypotheses, and estimate parameters within specified demographic models. A wide range of tools is available, utilizing either backward or forward simulations, with varying levels of support for parameter estimation.

Current coalescent-based simulators, such as `msprime`, generate stochastic tree sequences, offering significant flexibility while closely reflecting the data generation process (Baumdicker et al., 2021). However, their stochastic nature necessitates numerous replicates to obtain reliable estimates of summary statistics, especially when dealing with complex demographic models or statistics involving higher-order moments (Excoffier et al., 2013). Additionally, this inherent stochasticity complicates the use of gradient-based parameter estimation methods, often leading to the reliance on approximate Bayesian computation (ABC) (Kelleher and Lohse, 2020).

Another widely used category of tools is forward simulators, which offer the notable advantage of being able to incorporate selection (Gutenkunst et al., 2009; Jouganous et al., 2017; Haller and Messer, 2023). However, forward simulators have their own caveats—notably determining an optimal runtime to reach equilibrium—and demographic models are often easier to specify in a coalescent framework. Tools such as `dadi` and `moments`, rely on diffusion-based computations and provide exact solutions for a variety of summary statistics (Gutenkunst et al., 2009; Jouganous et al., 2017). Yet, closed-form solutions to the underlying differential equations are generally unavailable, requiring numerical approximations that can cause instabilities and complicate the computation of higher-order moments (Jouganous et al., 2017). `SLiM`, by contrast, explicitly tracks each individual in the population, offering high customizability and full access to evolutionary history (Haller and Messer, 2023). This level of detail, however, comes at the cost of increased computational complexity, which may be unnecessary for analyses focused solely on summary statistics. Moreover, as with `msprime, SLiM` simulations are stochastic and require many replicates for robust summary statistics estimation.

Phase-type theory presents a unified framework for modeling mixtures and convolutions of exponential distributions (Bladt and Nielsen, 2017). It offers a complementary method to the above approaches by providing exact, numerically stable solutions for a range of coalescent tree statistics (Hobolth et al., 2024). Phase-type theory models the coalescent process as a continuous-time Markov chain, with states representing the possible configurations of lineages in the population. Several software packages have been developed to support its application in population genetics. `PhaseTypeR` provides a general framework for utilizing phase-type distributions, with a focus on population genetics applications (Rivas-González et al., 2023). In `ptdalgorithms`, the emphasis is on accelerating computations by employing a graph-based approach (Røikjer et al., 2022). However, these tools typically require a solid understanding of the underlying theory, with state space construction often left to the user. Additionally, they offer only limited support for time-inhomogeneous coalescent processes, such as those involving changing population sizes or migration rates. Recently, Wences et al. (2024) demonstrated how to derive the distribution and moments of the TMRCA under various time-inhomogeneous coalescent models using phase-type theory. However, available quantities are limited to the TMRCA, and no software implementation is provided.

Here, we introduce `PhaseGen`, which supports the computation of branch length moments under piecewise-constant demographies. Branch length moments include quantities such as the expected length, variance, and correlation between the lengths of specific branches in a coalescent tree, and are key determinants of the patterns of polymorphism observed in genetic data. `PhaseGen` is particularly well-suited for optimizing demographic models in a coalescent framework, especially when multiple-merger coalescents (MMCs) are involved. It is also valuable for new methods that rely on exact coalescent-based solutions, making it unnecessary to rederive these quantities. The framework is also well-suited for exploratory analyses, where rapid model specification and easy access to a wide range of summary statistics are beneficial.

The paper is structured as follows: Section 2 begins with an overview of `PhaseGen`’s implementation, followed by computation and parameter estimation examples that demonstrate its capabilities. Section 3 then discusses potential future applications and perspectives on phase-type theory and software in population genetics. Additionally, Appendix A presents linear algebraic computation examples, followed by the theoretical framework of time-inhomogeneous phase-type distributions. The remainder of the appendix covers the state space and transition rates for various coalescent models (Appendix B), additional code examples (Appendix C), state space sizes (Appendix D), computation times under different demographic scenarios (Appendix E), and a summary of software validation against `msprime` (Appendix F).

## 2 Applications of PhaseGen

### 2.1 Software Implementation

`PhaseGen` is Python-based, although an R interface with usage examples is also available. It works internally by constructing the state space for the specified coalescent scenario, along with the weights associated with each state needed to compute the summary statistic of interest (Appendix Section A). `PhaseGen` enables the computation of branch length (cross-)moments of arbitrary order under a variety of coalescent models (`Kingman, Beta`, and `Dirac` coalescent) and demographic scenarios. Alternatively, custom weights (i.e., `Rewards`) can be assigned to each state in the state space, allowing for the computation of targeted quantities such as the time spent in a particular deme (cf. phasegen.readthedocs.io/en/v1.0.1/reference/Python/rewards.html). Additional features include access to the cumulative distribution function (CDF), probability density function (PDF), and quantile function for the coalescent tree height. Branch length moments can also be computed over a specified time interval by adjusting start and end times. By progressively increasing the end time, the accumulation of moments over time can also be analyzed.

The software package is equipped with a `Demography` interface akin to `msprime`, designed to handle temporal changes, with helper functions available for discretizing continuous demographic functions. To ensure correctness, `PhaseGen` has been extensively tested against stochastic estimates from `msprime`, comparing various summary statistics across a wide range of demographic scenarios, with no discrepancies observed (Appendix F). The source code is hosted on GitHub (github.com/Sendrowski/PhaseGen), with comprehensive documentation available at `phasegen.readthedocs.io`. Installation is facilitated via the pip package manager.

`In PhaseGen`, summary statistics under a given demographic model are obtained by creating a `Coalescent` object—the main entry point for all computations. This object encapsulates the distributional properties of the underlying coalescent process and includes the specified demography (`Demography`), coalescent model (`CoalescentModel`), lineage configuration (`LineageConfig`), locus configuration (`LocusConfig`), and the resulting state space (`StateSpace`). The `Demography` consists of one or more `Epochs`, within which population sizes and migration rates are constant. Internally, `PhaseGen` constructs a state space representing the possible `States` of the coalescent process (cf. Appendix A). For each epoch, the `Transition` rates between states are computed based on the specified coalescent model and demography. For tree height and total branch length statistics, the `LineageCountingStateSpace` is used, which tracks the number of lineages over time. For SFS-based statistics, the larger `BlockCountingStateSpace` is required, which tracks the number of branches subtending *i* lineages in the coalescent tree. `LineageConfig` and `LocusConfig` specify the initial number of lineages in each deme and locus, respectively. Upon creation, the `Coalescent` object provides access to a variety of statistics, the most commonly used of which are available as cached properties that are evaluated lazily–meaning they are computed only upon access and reused thereafter. This includes branch length moments of the tree height (`TreeHeightDistribution`), total branch length (`total branch length`), and the site frequency spectrum (`UnfoldedSFSDistribution` and `FoldedSFSDistribution`). In addition, `TreeHeightDistribution` supports computing the `CDF, PDF`, and `quantile` functions. For more complex statistics, `Rewards` can be used to assign weights to states in the state space and passed to the `Coalescent.moment` function. For further details, please refer to the online reference, accessible through the monospaced terms linked in the text.

In this section, we showcase the capabilities of `PhaseGen`. We begin with the computation of summary statistics for the Kingman coalescent (Section 2.2) and progressing to more complex scenarios, including piecewise-constant demographies (Section 2.3), migration (Section 2.4), multiple-merger coalescents (Section 2.5), and recombination (Section 2.6). We then demonstrate the utility of `PhaseGen` for parameter estimation (Section 2.7), and show how the inclusion of higher-dimensional summary statistics can improve inference accuracy (Section 2.8).

### 2.2 Standard Coalescent

In this example, we define a Kingman coalescent model with *n* = 8 lineages, and a constant population size of *N* = 1 (Wakeley, 2009). Note that in most examples we use a population size of 1 for brevity, since coalescent rates scale linearly with *N*. Figure 1 displays the tree height density, expected SFS, and 2-SFS correlation matrix, along with the code used to define the model and compute these statistics. We create a `Coalescent` object using the standard coalescent model with a single population of size 1, both of which are default settings. The tree height density (`coal.tree height.pdf`), expected SFS (`coal.sfs.mean`), and 2-SFS correlation matrix (`coal.sfs.corr`) are then directly accessible. See Section 2.8 for a more detailed introduction to the 2-SFS. Note that while these estimates are based on branch lengths, they can be scaled by the mutation rate to derive the expected number of observed mutations under the infinite-sites model.

**Figure 1:**
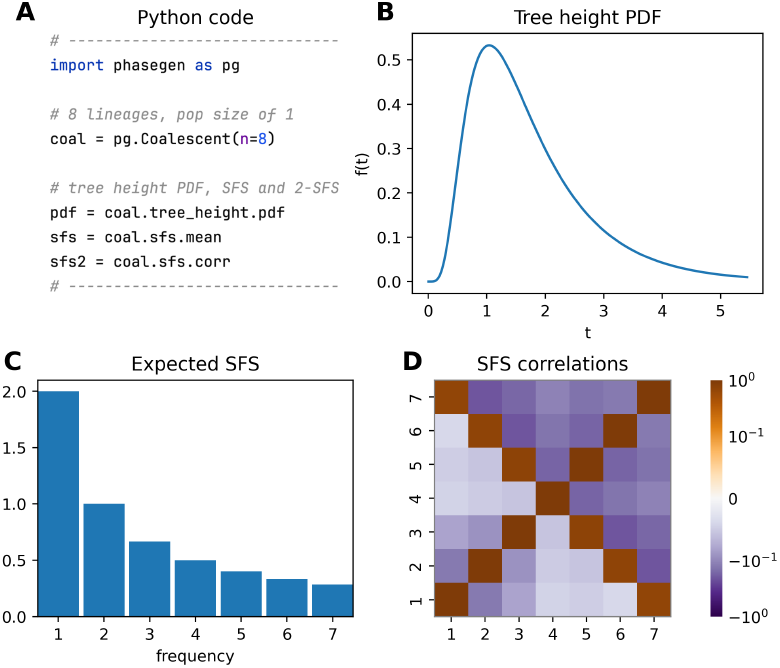
Kingman coalescent model with *n* = 8 lineages and constant population size *N* = 1. **A**: Python code snippet defining the coalescent object for this scenario, and accessing the tree height density, expected SFS, and 2-SFS correlation matrix (cf. plot_manuscript_kingman_example.py, containing the full code to generate the figure). **B**: Tree height density (PDF). **C**: Expected SFS. **D**: (symmetric) 2-SFS correlation matrix representing the branch length correlation of branches subtending i and j lineages in the coalescent tree.

### 2.3 Multiple Epochs

Piecewise-constant demographic models can be specified using a Python dictionary that maps time points to population sizes. In this example, we define a 2-epoch model where the population size decreases backward in time from *N* = 1 to 0.2 at *t* = 1. Figure 2 shows the tree height density, expected SFS, and 2-SFS correlation matrix for this model, along with the code used to define the `Coalescent` object. The population size decrease leads to a higher coalescent rate at time *t* = 1, as reflected in the tree height density. The expected SFS shows a relative excess of singleton variants, due to relatively longer terminal branches resulting from the initially larger population size.

**Figure 2:**
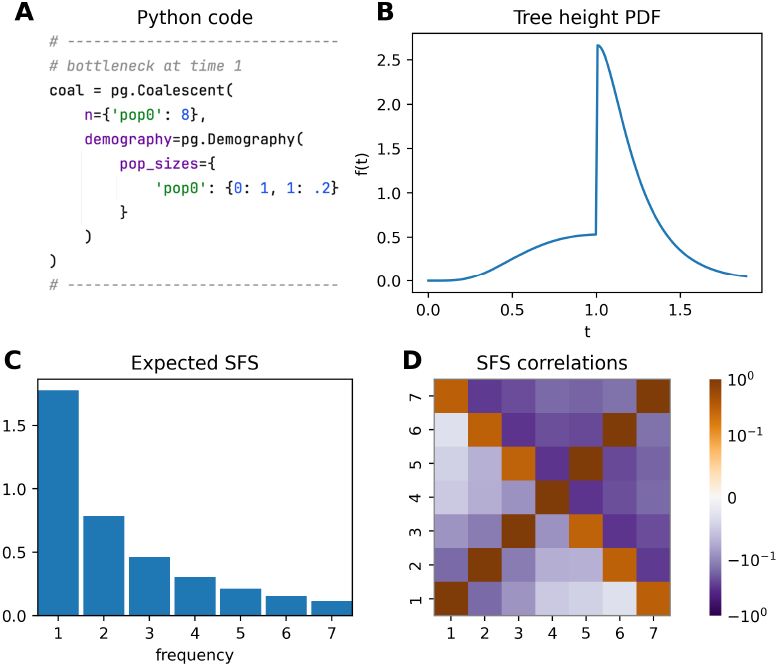
Two-epoch demography featuring a population size reduction backward in time. **A**: Python code snippet defining the coalescent object for this scenario (cf. plot_manuscript_2_epoch_example.py, containing the full code to generate the figure). The dictionary defines the population sizes, where keys represent time points, and values specify the corresponding population sizes. The identifier pop0 is used for the single population. **B**: Tree height density (PDF), highlighting a sudden increase in the coalescent rate due to the population size reduction. **C**: Expected SFS, showing a relative deficit of high-frequency variants. **D**: 2-SFS correlation matrix.

### 2.4 Isolation with Migration

We can also define demes and specify migration between them (Hey, 2009). In this example, we model two demes, `pop0` and `pop1`, with population sizes 1 and 2, respectively (Figure 3). Migration occurs backwards in time at a per-generation rate of *m* = 0.3 from `pop0` to `pop1`. Both demes begin with *n* = 4 lineages in the present. Here, we also marginalize the SFS over demes, i.e., we weight the branch lengths by the proportion of total lineages spent in each deme over time. In this example, branches are longer for lineages in deme `pop1` due to its larger population size and thus lower coalescent rate. Also note that the resulting SFS shows an excess of quadrupletons, which corresponds to relatively longer branches subtending four lineages in the average coalescent tree. This pattern arises because coalescence tends to occur within demes before migration.

**Figure 3:**
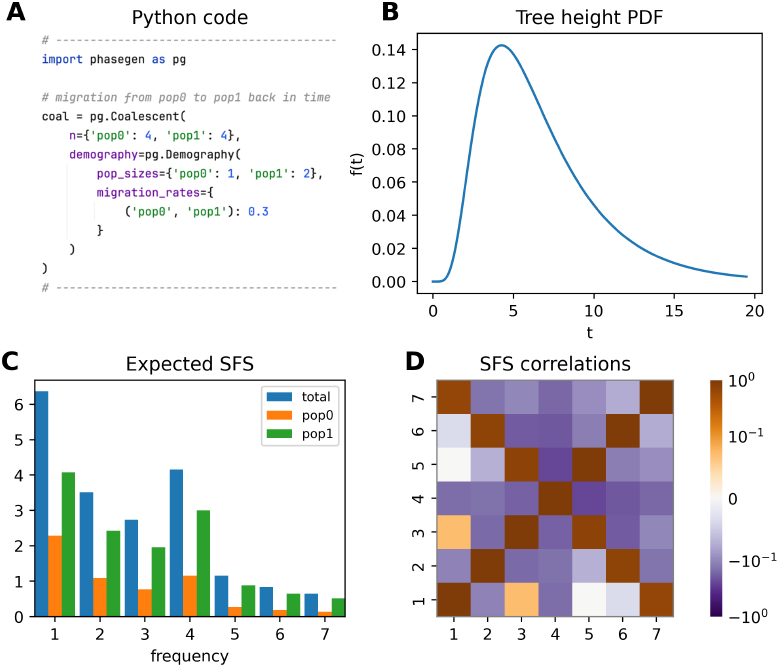
Two-deme model with uni-directional migration. **A**: Python code snippet defining the coalescent object for this scenario (cf. plot_manuscript_migration_example.py, containing the full code to generate the figure). The first element of the migration rate tuple specifies the source deme (`pop0`), and the second specifies the target deme (`pop1`) backwards in time. **B**: Tree height density (PDF). **C**: Total expected SFS and deme-specific contributions corresponding to the proportion of total lineages spent in each deme over time for branches subtending a given number of lineages in the coalescent tree. **D**: 2-SFS correlation matrix, revealing a positive correlation between singletons and tripletons in the SFS. Note that the 2-SFS is not to be confused with the joint-SFS, which records the co-occurrence of allele frequency counts across demes.

### 2.5 Multiple-Merger Coalescent

PhaseGen also supports multiple-merger coalescents (MMCs), where more than two lineages can coalesce simultaneously (see Section 2.8 for a more detailed introduction to MMCs). Here, we use illustrate the `BetaCoalescent` model, where the probability of coalescence among a fraction of lineages is determined by a beta distribution parameterized by *α*, ranging from 1 (star-like coalescent) to 2 (Kingman coalescent) (Schweinsberg, 2003). In this example, *α* = 1.4, leading to frequent coalescence events involving more than two lineages (Figure 4). The resulting expected SFS shows a relative excess of high-frequency variants, and the 2-SFS correlation matrix reveals positive correlations between branches of different frequency bins—both typical signatures of MMCs.

**Figure 4:**
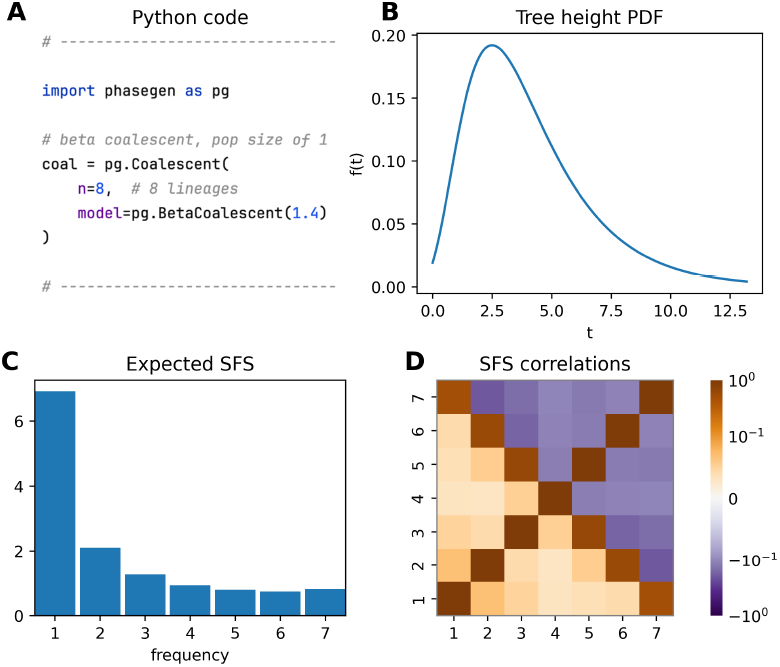
Beta coalescent model with *α* = 1.4, *n* = 8 lineages, and constant population size *N* = 1. **A**: Python code snippet defining the coalescent object for this scenario (cf. plot_manuscript_mmc_example.py, containing the full code to generate the figure). **B**: tree height density (PDF). Note that unlike for the Kingman coalescent, the TMRCA may be very close to zero. **C**: expected SFS which is characterized by a relative deficit of doubletons and an excess of high-frequency variants. **D**: 2-SFS correlation matrix showing a positive correlation between branches of different frequency bins.

All dynamics presented so far can be combined into a single model (see Appendix C.1 for a more complex example).

### 2.6 Coalescent with Recombination

There exist simple closed-form solutions for the covariance of the TMRCAs between two loci depending on the recombination rate *ρ*. Specifically, for two lineages, the covariance is given by

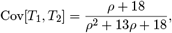

where both loci are assumed to be initially linked (cf. Formula 7.28 in Wakeley (2009)). Here, *ρ* = 4*Nr* is the scaled recombination rate, with *r* denoting the per-generation probability of recombination between the two loci.

To explore this further, we may wish to compute the tree height covariance under a timeinhomogeneous demographic model. Figure 5 illustrates the covariance under a 2-epoch model where the population size changes from 1 to *N*_1_ at time *t* = 1. For a fixed *ρ*, larger population sizes lead to a higher covariance due to the linear increase in TMRCA with population size. However, the correlation decreases with increasing *ρ* because higher recombination rates allow more opportunities for recombination events to occur. Notably, this relationship is not significantly altered by the time-inhomogeneous demography. Note that SFS-based summary statistics are currently not supported for multiple loci, since this would require a more complex state space (cf. Section 3 for a discussion on this topic).

**Figure 5:**
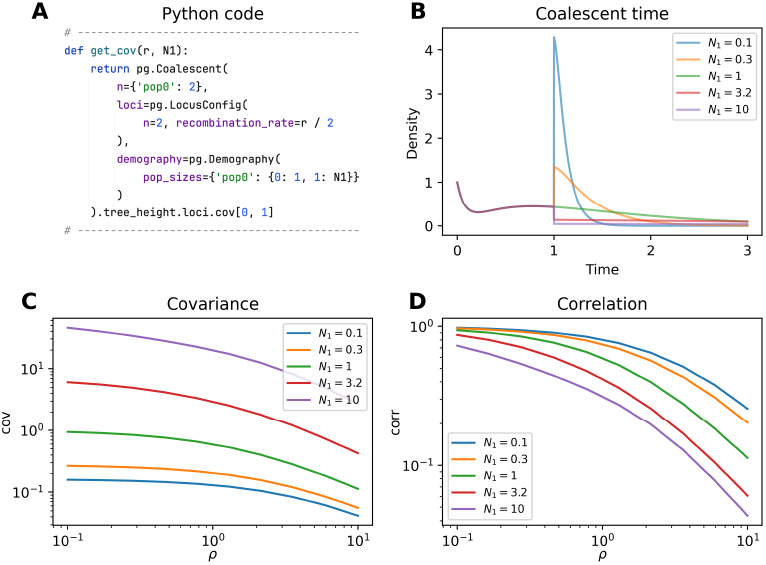
2-epoch demography with a population size change from 1 to *N*_1_ at time *t* = 1. **A**: Python function to compute the covariance of the TMRCAs depending on *ρ* and *N*_1_ (cf. plot_manuscript_recombination_example.py, containing the full code to generate the figure). Note that *ρ* is scaled differently in `PhaseGen` so that we divide by 2 here. **B**: tree height densities for the total TMRCA of both loci for different values of *N*_1_ and *ρ* = 10. Note the initial slump in density which is due to recombination occurring before coalescence. **C**: Covariance of the TMRCAs for different values of *ρ* and *N*_1_. **D**: Correlation of the TMRCAs, which ranges from 1 for low *ρ* to 0 for high *ρ*. Note that the covariance increases with *N*_1_, while the correlation decreases.

### 2.7 Statistical Inference

We begin by establishing a connection between the SFS computable from real data and the SFS provided by `PhaseGen`. The site frequency spectrum (SFS) is a summary statistic that records the frequencies of alleles at sites in a sample of *n* individuals. That is, each entry *k* in the SFS counts the number of sites where the derived allele appears in exactly *k* individuals. Under the infinitesites model, each site is assumed to experience at most one mutation, so the frequency of a derived allele at a site is directly related to the structure of the underlying coalescent tree—specifically, to the number of terminal branches subtended by the branch where the mutation occurred. The coalescent tree is shaped by selection and demography, but also reflects random variation from the stochastic nature of the coalescent process. By averaging over many sites, and assuming sufficient recombination, we reduce this stochastic noise by effectively sampling many independent realizations of the coalescent process. `PhaseGen` computes the expected SFS under a specified demographic model, with each frequency bin equal to the total branch length where mutations produce alleles at that frequency. Assuming a constant mutation rate and selective neutrality, mutations follow a Poisson process, so that the expected number of mutations at frequency *k* is given by *θ* times the total branch length subtending *k* lineages, where *θ* is the population-scaled mutation rate. Thus, multiplying the expected SFS from `PhaseGen` by *θ* gives the expected SFS observable in real data.

The availability of exact solutions for summary statistics like the SFS lends itself to gradientbased parameter estimation, and thus, `PhaseGen` provides a compact framework for this purpose. Gradient-based optimization relies on computing the derivative of the loss function with respect to model parameters, enabling navigation toward optimal values via gradient descent. The gradient is often computed numerically because the loss function is usually too complex to differentiate analytically.

Within `PhaseGen`, parameter inference under a given demographic model can be performed using the `Inference` class. This involves defining a function that returns a parameterized `Coalescent` object (`coal`). In each optimization iteration, this function is called to obtain the corresponding `Coalescent` object for the current parameter values. In this example, we model a single population size change from size 1 to `Ne` at time `t`, where both the time of change `t` and the resulting population size `Ne` are variable. The analysis is based on an unfolded SFS obtained from a whole-exome Scandinavian silver birch dataset (Salojarvi et al., 2017; Sendrowski, 2022) (`observation`). Ancestral states were determined using two outgroup elder species (Alnus incana & A. glutinosa). To evaluate model fit during optimization, we must specify a loss function (loss) which takes two arguments: the observed summary statistic and the parameterized `Coalescent` object. In this example, the loss function is a composite Poisson likelihood, which assumes that mutations at different frequencies are independent Poisson variables (Poisson random field assumption). To standardize the data, the SFS is normalized by Watterson’s estimator of the population-scaled mutation rate *θ*. Note that we multiply *θ* by the total number of sites in the loss function because *θ* is computed on a per-site basis. By default, 10 independent optimization runs are carried out with different initial values, and the best result is selected. Optionally, a resampling function can be provided to perform parametric bootstrapping (`resample`). By default, 100 bootstrap replicates are performed, each initialized with the best fit from the original optimization run. Parameter bounds are set to [0, 4] for t and [0.1, 10] for Ne (`bounds`). Note that both `t` and `Ne` are expressed in units relative to the effective population size, as both scale linearly with it. The absolute values can be obtained by multiplying the estimates by the effective population size. The inference results are visualized in Figure 6, and indicate a population size reduction in the past (cf. plot_manuscript_inference_example.py). The inference framework permits an arbitrary parameterization of the coalescent distribution and a customizable choice of loss function, and is thus rather flexible. Additional inference features include support for distributed computing on clusters (cf. [the package documentation](https://phasegen.readthedocs.io/en/v1.0.1/reference/Python/inference.html) for more details).

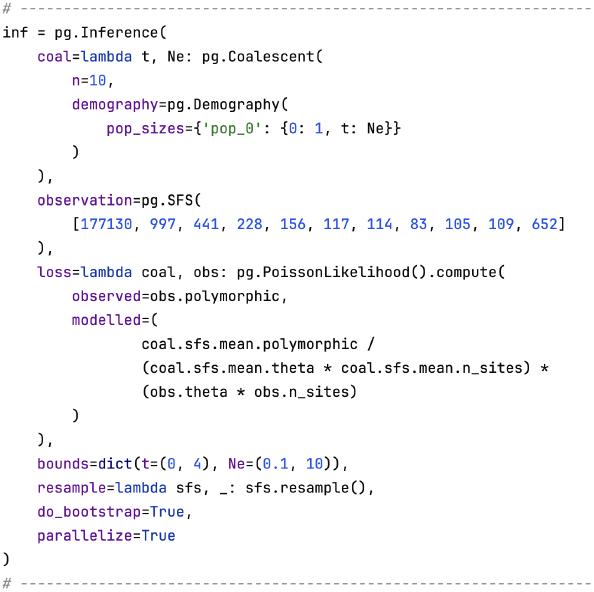

**Figure 6:**
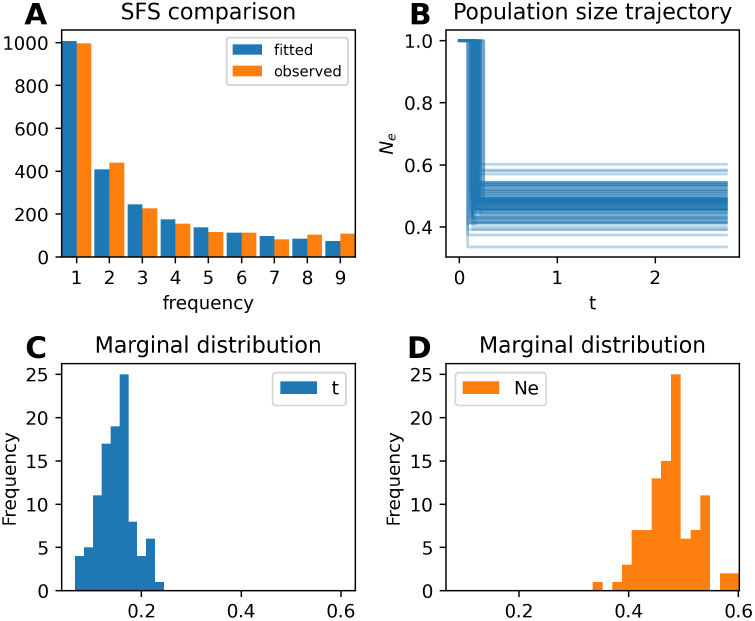
**A**: Comparison between the fitted and observed SFS. **B**: Population size trajectories derived from the initial parameter estimate and bootstrap replicates, indicating a population size reduction in the past. **C & D**: Marginal distributions of bootstrap parameter estimates. The initial optimization took 3.88 seconds with 125 function evaluations per run, while each bootstrap run averaged 1.06 seconds and 32 evaluations on a MacBook Pro M2 (cf. Appendix E for a detailed discussion of computation times). Note that computations can be parallelized. For comparison, the same analysis using dadi took 0.71 seconds with 144 function evaluations per run, and each bootstrap run averaged 0.29 seconds and 146 evaluations (cf. run_2_epoch_inference example_dadi.py).

### 2.8 Inference Example on SFS Covariance

The following example demonstrates the advantages of including higher-dimensional summary statistics for parameter estimation, with a focus on the 2-SFS. As a two-dimensional extension of the SFS, the 2-SFS records the covariance between different frequency bins in the SFS, i.e. the branch length covariance of branches subtending *i* and *j* lineages in the coalescent tree. In particular, it can reveal signals of multiple merger coalescents (MMCs), which involve the simultaneous coalescence of more than two lineages (Birkner et al., 2013b). Such events may arise in populations with highly skewed offspring distributions (Eldon and Wakeley, 2006). MMCs generate a distinct positive correlation between different frequency bins of the SFS—a signal that has been shown to be robust to confounding factors such as demographic changes and recombination (Fenton et al., 2025). In practice, the 2-SFS can be computed from data by considering the frequencies of nearby SNP pairs, assuming negligible recombination between them. This requires careful consideration of the mutational opportunities, after which the raw 2-SFS can be centered and normalized to obtain a correlation matrix. Alternatively, inference can be performed directly on the uncentered 2-SFS (see Fenton et al. (2025) for alternative summary statistics and further discussion).

To illustrate the value of incorporating higher-order moments in inference, we present a proof-of-concept example using `PhaseGen`, comparing results from SFS-only inference to those using both the SFS and the 2-SFS correlation matrix (Figure 7). The ground truth in this example assumes a 3-epoch demographic model with a temporary increase followed by a drastic decrease in population size backward in time, and a beta coalescent with *α* = 1.7, making multiple mergers relatively frequent. More precisely, the population size changes from *N* = 1 to 4.5 at *t* = 0.5, and subsequently to 0.1 at *t* = 4.5. The demographic scenario was specifically chosen to generate an SFS lacking the typical U-shaped profile associated with multiple mergers, making it difficult to infer *α* from the SFS alone. For inference, a simplified 2-epoch demography is fitted, estimating a population size change from size 1 to *N*_1_ at time *t*_1_, alongside *α*. Note that different models are used for inference and ground truth, introducing noise and reflecting the fact that, in real data, the actual demographic history is more complex than the inference model.

**Figure 7:**
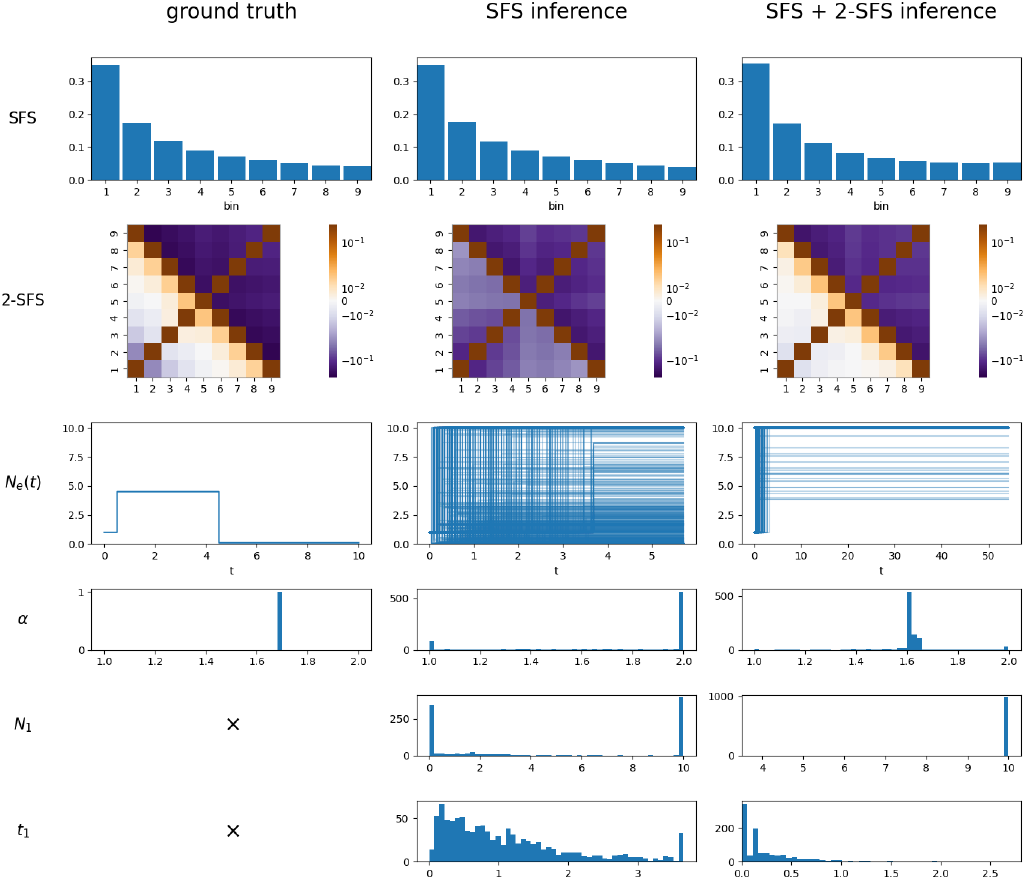
Inference results for Section 2.8, showing ground truth (left), SFS-based inference (middle), and SFS+2-SFS-based inference (right). The rows represent the SFS, 2-SFS correlation matrix, population size trajectory, and inferred values for *α, N*_1_, and *t*_1_, respectively. The ground truth demography includes a population size increase from *N* = 1 to 4.5 at time 0.5, followed by a sharp decline to 0.1 at time 4.5. The demography is challenging to infer from the SFS alone, leading to substantial variability in both estimates of *α* and the inferred population size history (cf. infer_mmc_kingman_2sfs.py, plot_mmc_inference.py). For the inference procedure including the 2-SFS, the initial optimization took 29.93 seconds per run, and each bootstrap run averaged 13.94 seconds on a MacBook Pro M2. Note that computations can be parallelized.

A multinomial likelihood is applied to the SFS, which is normalized by the total number of polymorphic sites. When incorporating the 2-SFS, an additional loss term is added, describing the absolute difference between the observed and modeled correlation coefficient for the 4th and 5th frequency bins. Using only a single 2-SFS coefficient was motivated by performance considerations, and this specific entry was found to be particularly informative. Bootstrapping is performed by applying Poisson sampling to the observed SFS data, generating 1,000 replicates. Figure 7 demonstrates that *α* is recovered reasonably well when augmenting the loss function with information from the 2-SFS, even though different demographic models are used for inference and ground truth. In contrast, SFS-only inference fails to recover the true value of *α*, with both *α* and the inferred population size trajectory showing significant variation across bootstraps. This is because the SFS is also sensitive to other demographic effects, making it challenging to disentangle the signal of multiple mergers from the effects of population size changes.

## 3 Discussion

In this work, we introduced `PhaseGen`, a software package providing an accessible interface for computing exact and numerically stable moments of piecewise-constant coalescent processes under various demographic scenarios. Such discrete rate changes are common in population genetics, and can approximate continuous demographies with arbitrary precision.

This framework utilizes Van Loan’s method for computing moments (Appendix A.8), which relies on matrix exponentiation, and for which accurate solutions can be obtained using Padé approximants (Van Loan, 1978; Al-Mohy and Higham, 2010). Matrix exponentiation generally has a cubic runtime complexity, depending on the sparsity and structure of the matrix. Consequently, it is crucial to keep the state space as small as possible; see Appendix D for state space sizes and Appendix E for corresponding benchmarking results across different configurations.

A graph-based approach, such as that implemented in `pdtalgorithms`, can significantly accelerate computations (Røikjer et al., 2022). However, computing time-inhomogeneous moments of order larger than one (e.g., second-order moments like variance) requires accounting for cross-moments between different epochs, which the graph-based approach does not currently support. Alternatively, faster matrix exponentiation algorithms, particularly for sparse matrices or those leveraging the structure of the Van Loan matrix, could offer substantial improvements. While GPU acceleration remains limited, it also holds potential for significant speedups in the future. Incorporating exact gradient information for parameter estimation is another promising approach that could substantially accelerate optimization algorithms, especially in high-dimensional parameter spaces. Future research could also explore dynamic state space reduction by merging equivalent states or collapsing states with negligible expected sojourn times, depending on the demographic model.

On a related note, the theory of probability generating functions in population genetics (Lohse et al., 2011) is closely tied to phase-type theory (Hobolth et al., 2024). The `agemo` software package, for instance, utilizes a graph-based approach to compute probabilities for small-sample-size SFS under coalescent-with-migration models (Bisschop, 2022). These mutational block configuration probabilities are particularly useful for fitting models to small datasets. `PhaseGen` also provides phase-type-based support for this, as detailed in Hobolth et al. (2025). However, its current framework is restricted to time-homogeneous demographies, although time-homogenous dynamics such as migration between demes, and different coalescent models are supported (cf. `the package documentation` for more details). Expanding its capabilities to include time-inhomogeneous scenarios would greatly enhance its applicability for modeling small datasets.

Future extensions to `PhaseGen` could include support for additional structured coalescents, such as models incorporating seed banks, diploidy, and polyploidy. The availability of full distribution functions for rewarded summary statistics, such as the total branch length or the SFS, would also be beneficial. However, to date, this does not appear to be generally possible for time-inhomogeneous models within the phase-type framework. Adding support for SFS-based statistics under the twolocus model would also be valuable, though this would necessitate lineage labeling, leading to a significant increase in state space size, thus making computations infeasible even for moderate lineage counts. Similarly, joint-SFS statistics, which record the co-occurrence of allele frequency counts across different demes, require lineage labeling, i.e. the deme of origin. Finally, a fully labeled state space implementation could enable the calculation of many additional statistics, but computational feasibility would remain a challenge for more than 6–7 lineages, depending on the number of unique labels involved.

`PhaseGen` can also be used to compute summary statistics under the multi-species coalescent (MSC) model, i.e., by treating each species as a separate deme. Recently, Guerra and Nielsen (2022) derived the covariance of pairwise differences under the MSC model under a piecewise-constant demography, and examined the biases introduced in *F*_*ST*_ estimates derived from pairwise differences. While computing the covariance of pairwise differences requires additional lineage labeling, metrics like *F*_*ST*_ can be expressed as rational functions of pairwise coalescent times, making them accessible within `PhaseGen` (Slatkin, 1991).

In conclusion, we hope that the software and theory presented here will serve as valuable resources for the population genetics community, facilitating exploratory analyses and the fitting of demographic models to data.

## Acknowledgements

The authors would like to thank Mogens Bladt for helpful discussions, as well as the reviewers for their valuable comments and suggestions. This work has been supported by the Novo Nordisk Foundation (Data Science Collaborative Research Programme, grant 0069105).

## Data Availability

The data underlying this article are available on Zenodo, at https://doi.org/10.5281/zenodo.14880470.

## Appendix A Theory

This section provides the theoretical background on time-inhomogeneous phase-type distributions, which form the basis of `PhaseGen`. It begins with two illustrative computation examples designed to build intuition while minimizing theoretical complexity, followed by a detailed explanation of the underlying theory.

### A.1 One-Epoch Computation Example

As a starting example, we compute the expected length of singleton branches (𝔼[*L*_1_]) under the standard Kingman coalescent model with a constant population size. Singletons are branches in the coalescent tree that subtend a single terminal lineage, so that, under the infinite-sites assumption, their lengths determine the probability of observing singleton mutations in a population (Figure A1). We later extend this example to a more complex case involving a 2-epoch demography and the covariance of singleton and doubleton branch lengths (Section A.2).

Assume we have *n* = 3 lineages in a single deme with constant population size *N* = 1. This gives rise to a state space of 3 states {*e*_3_, *e*_2_, *e*_1_}, corresponding to the number of lineages present at a given time. From Figure A1, it is apparent that we have 3 singleton branches in state *e*_3_ and 1 in state *e*_2_. The expected total singleton branch length *L*_1_ is given by the number of singleton branches in each state multiplied by the expected time spent in that state, i.e., 𝔼[*L*_1_] = 3𝔼[*T*_3_] +𝔼[*T*_2_], where *T*_*i*_ denotes the time spent in state *e*_*i*_. In the Kingman coalescent model, the time spent in each state is exponentially distributed with rate 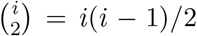, where *i* is the number of lineages. Each transition reduces the number of lineages by 1 until the absorbing state *e*_1_ is reached. We can summarize the transition rates in the following rate matrix:

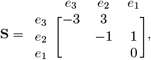

where off-diagonal zero entries have been left blank for brevity. Note that diagonal entries are chosen such that each row sums to zero.

We now define a reward vector which records the number of singletons in each state. We have *r* = (3, 1, 0), which can be summarized in the diagonal reward matrix:

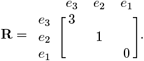

To compute the expected singleton branch length, we use Van Loan’s method (cf. Section A.8), which involves constructing the 2 *×* 2 block matrix containing the rate and reward matrices. We

**Figure A1:**
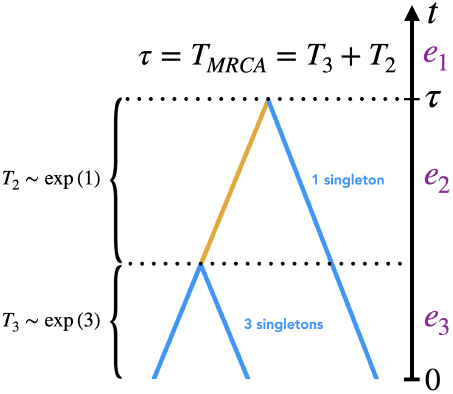
Schematic of a standard Kingman coalescent tree with *n* = 3 lineages under constant population size of *N* = 1. The waiting time in each state is exponentially distributed with rate 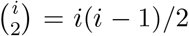, where *i* is the number of lineages. There are 3 singleton branches in state *e*_3_ and 1 in state *e*_2_. State *e*_1_ is absorbing, with *T*_*i*_ denoting the time spent in state *e*_*i*_. The time to the most recent common ancestor, *T*_*MRCA*_, equals the absorption time *τ*. The total singleton branch length *L*_1_ is given by 3*T*_3_ + *T*_2_.

have

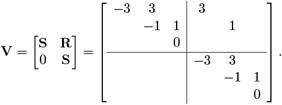

We then compute the matrix exponential in the limit as time approaches infinity to ensure the process reaches the absorbing state, i.e., all lineages have coalesced:

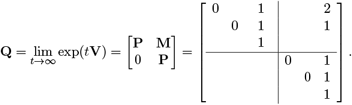

From **Q**, we extract

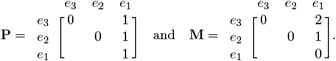

Matrix **P** gives the probability of ending in each state, conditioned on the starting state. We observe that the process ends in state *e*_1_ with probability 1, regardless of the starting state. Matrix **M** contains the singleton branch lengths with respect to starting and ending states. Specifically, the branch length is **M**(*e*_3_, *e*_1_) = 2 when starting from *e*_3_ and ending in *e*_1_, and **M**(*e*_2_, *e*_1_) = 1 when starting from *e*_2_ and ending in *e*_1_. Van Loan’s method also allows for the computation of higher-order moments by creating larger block matrices, and piece-wise constant demographies can be handled by multiplying the block matrices for each epoch.

Let ***α*** = (1, 0, 0) be the initial state distribution (since we start with 3 lineages with a probability of 1), and let **e** = (1, 1, 1)^*⊤*^ be the exit vector summing up the branch lengths for each end state. Then, the expected singleton branch length is given by

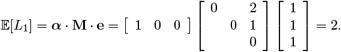

Alternative, we could have directly computed 𝔼[*L*_1_] = 3 *·* 𝔼[*T*_3_] + 𝔼[*T*_2_] = 3 *·* 1(*/*3)+ 1 = 2, using 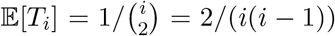, the mean of an exponential distribution with rate 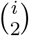. However, the matrix machinery introduced here generalizes naturally to more complex scenarios, such as those presented in the next section.

#### A.2 Two-Epoch Computation Example

Here, we compute the covariance of singleton and doubleton branch lengths under a 2-epoch demography using time-inhomogeneous phase-type theory. Singletons and doubletons are branches in the coalescent tree that subtend one and two lineages, respectively, so that their lengths determine the number of mutations observed in a population at certain frequencies (Figure A2). We want to determine Cov[*L*_1_, *L*_2_] = 𝔼[*L*_1_*L*_2_] *™* 𝔼[*L*_1_]𝔼[*L*_2_], where *L*_1_ and *L*_2_ are random variables representing singleton and doubleton branch lengths, respectively. Assume we have *n* = 3 lineages in a single deme, with a population size of *N*_1_ = 1 for *t ∈* [0, 2) and *N*_2_ = 0.5 for *t ∈* [2, *∞*). This gives rise to a state space of 3 states {*e*_1_, *e*_2_, *e*_3_} corresponding to the number of lineages present a(t a) given time. The transition rates between states are determined by the coalescent rate, which is 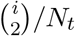 where *i* is the number of lineages and *N*_*t*_ the population size at time *t*. We can summarize the transition rates by constructing rate matrices for the two epochs. We have

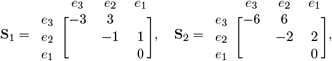

where off-diagonal zero entries have been left blank for brevity.

Next, we define reward vectors which describe the number of single- and doubleton branches in each state. Rewards assign a weight to each state, which is chosen depending on the summary statistic of interest (cf. Section A.5). Here we define *r*_1_ = (3, 1, 0) and *r*_2_ = (0, 1, 0) for single- and doubleton branch lengths, respectively (Figure A2). From this we construct the diagonal reward matrices **R**_*i*_ = △(*r*_*i*_). We have

**Figure A2:**
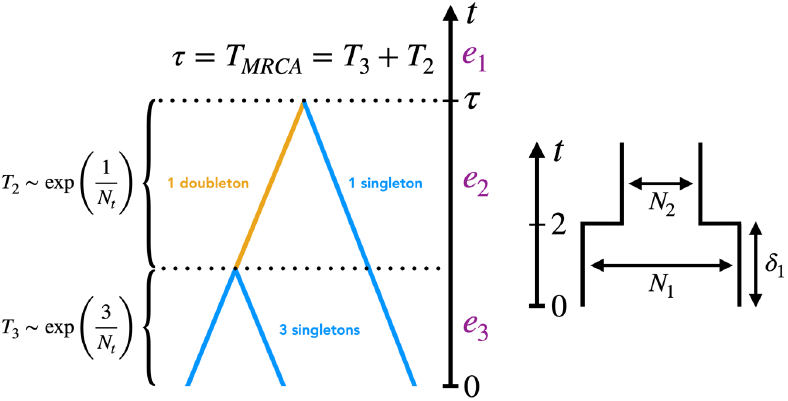
Schematic of a standard Kingman coalescent tree with *n* = 3 lineages (left) under a 2-epoch demography (right). The rate of coalescence is given by 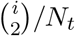 where *i* is the number of lineages, and *N*_*t*_ the population size at time *t*. If the population size were constant, *T*_2_ and *T*_3_ would be independent. For varying population sizes, however, the coalescent rate is time-dependent so that *T*_2_ depends on *T*_3_. We count 3 singleton and 0 doubleton branches in state *e*_3_, and 1 singleton and 1 doubleton branch in state *e*_2_. State *e*_1_ is absorbing, and *T*_*i*_ denotes the time spent in state *e*_*i*_. The time to the most recent common ancestor *T*_*MRCA*_ is given by the time to absorption *τ*.

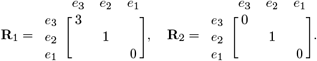

We first compute the uncentered branch length covariance (𝔼[*L*_1_*L*_2_]), and center it subsequently. To achieve this, we make use of Van Loan’s method (cf. Sections A.8 and A.9), which involves constructing a block matrix for each epoch containing the reward and rate matrices, which are then exponentiated. For epochs 1 and 2, respectively, we have the 3 *×* 3 block matrices

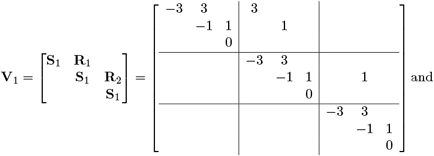

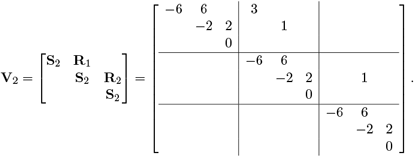

The accumulated reward per epoch can then be computed by exponentiating the block matrices over the time spent in each epoch. We obtain

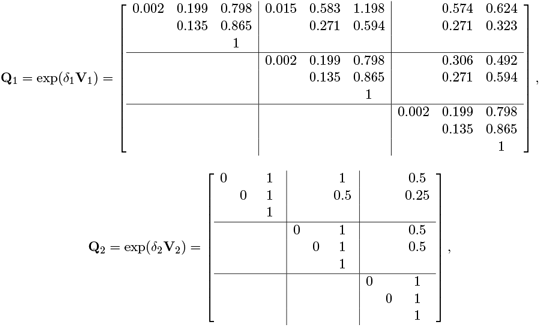

where *δ*_1_ = 2 and *δ*_2_ *→ ∞*, since the last epoch is infinite.

We now provide some interpretation for the quantities contained in **Q**_1_ and **Q**_2_. Let

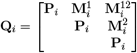

be the exponentiated block matrix for epoch *i*. The *jk*-th entry of **P**_*i*_ is the probability of being in state *k* at the end of epoch *i* given that the process started in state *j* at the beginning of epoch *i*. For example, let

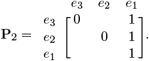

Since epoch 2 is infinite, {**P**_2_}_*i*3_ = 1, i.e., the process ends in the absorbing state *e*_1_ with probability ~ 1 for all starting states. Matrices 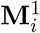 and 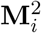 contain the accumulated reward in epoch *i* for single- and doubleton branches, respectively. Consider for instance 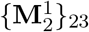, which contains the accumulated reward for singleton branches in epoch 2 given that we enter epoch 2 in state *e*_2_. The mean time spent in *e*_2_ is 1*/*2, during which there is exactly one singleton branch. Hence 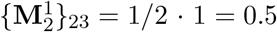. At last,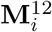can be interpreted as the accumulated uncentered covariance of single- and doubleton branches in epoch *i*, conditional on singletons occurring before doubletons.

To combine the results for the two epochs, we compute

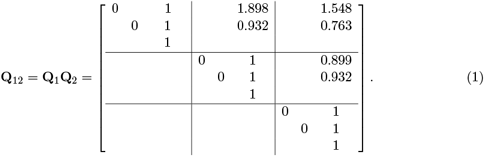

Note that by considering the individual blocks of **Q**_1_**Q**_2_, we obtain

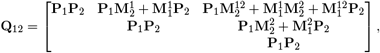

which shows how the epochs are combined for each block. Notably, the upper right block is 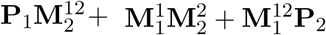, which effectively combines the accumulated covariance across epochs.

More precisely, let the upper right block of **Q**_12_ be

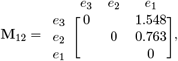

which contains the accumulated uncentered covariance conditioned on the order of single- and doubleton branches depending on start and end states (denoted as 𝔼[*L*_1_*L*_2_|**R**_1_, **R**_2_]).

We can now obtain our results with respect to the start and end states by

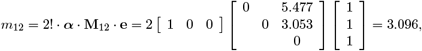

where ***α*** is the initial state probability vector which is (*e*_3_, *e*_2_, *e*_1_) = (1, 0, 0), as we start with 3 lineages with a probability of 1.

Since this result is conditioned on the order in which is the single- and doubleton rewards are accumulated (cf. Section A.9), we repeat the above procedure with the reward matrices permuted, from which we obtain *m*_21_ = 1.391. In addition, to center 𝔼[*L*_1_*L*_2_], we also need to obtain the mean single- and doubleton branch lengths (𝔼[*L*_1_] and 𝔼[*L*_2_]). We can do this by constructing and exponentiating the 2 *×* 2 block matrices

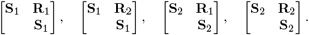

These matrices are already contained in the block matrices **V**_1_ and **V**_2_ above. Due to the nested structure of these block matrices, combined with matrix exponentiation, the mean branch length results are already contained in **Q**_12_. Consequently, we obtain a mean singleton branch length of

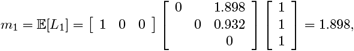

and, by a similar procedure, a mean doubleton branch length of *m*_2_ = 0.899.

Finally, we can compute the singleton/doubleton covariance by

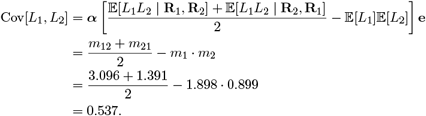

Figure A3 shows Cov[*L*_1_, *L*_2_] for different values of *N*_1_, *N*_2_, *δ*_1_ and *δ*_2_. As we will later see, this computation example can straightforwardly be generalized to cross-moments of arbitrary order and reward structure under more complex demographic scenarios.

**Figure A3:**
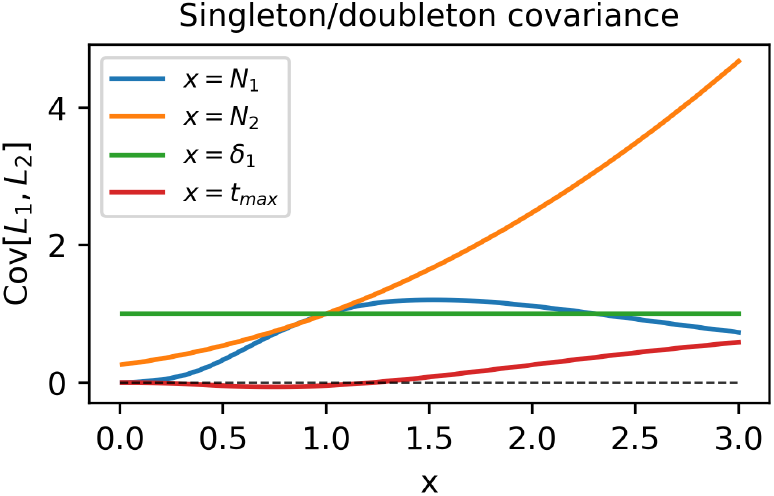
Singleton/doubleton branch length covariance for varying population sizes (*N*_1_, *N*_2_), time of population size change (*δ*_1_), and stopping time (*t* _max_). The remaining values were fixed to (*N*_1_, *N*_2_, *δ*_1_, *δ*_2_) *→* (1, 1, 2, *∞*) as in the example. Interestingly, the covariance reaches its mode around *N*_1_ = 1.5 before becoming smaller for larger values of *N*_1_. Also note that single- and doubletons initially covary negatively (cf. *x* = *t* _max_). This is because observing doubletons early on implies that the first coalescence event has already taken place which in turn reduces the time spend with singletons.

#### A.3 Phase-Type Distributions

Consider a Markov jump process {*X*_*t*_}_*t≥*0_ with finite state-space *E* = {1, 2, …, *k*} and let **S**(*t*) = {*s*(*t*)_*ij*_}_*i,j*=1,…,*k*_ be the rate matrix holding the exponential rates of jumping from state *i* to *j* at time *t*. Define the time until absorption as

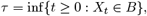

where *B ⊂ E* is the set of absorbing states (Albrecher and Bladt, 2018). The absorption time *τ* is said to be (time-inhomogeneously) phase-type distributed with state space *E*, initial state vector ***α*** = (*α*_*i*_)_*i∈E*_ and rate matrix **S**(*t*), and we write

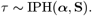

#### A.4 The Coalescent

We begin by defining the transition rates for the standard Kingman coalescent. Generalizations to more complex demographic scenarios involving multiple demes, multiple-merger coalescents and two loci are provided in the Appendix. We require a state space *E* = {1, …, *n*} with *n* states, where *n* denotes the number of initial lineages (Hobolth et al., 2019). The only absorbing state is *B* = {1}, and the transition rates are given by

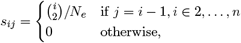

where *N*_*e*_ is the effective population size. Henceforth, we refer to the state space keeping track of the number of lineages at a time as the *lineage-counting state space* (see Appendix B.1.1 for the transition graph).

In order to obtain more complex summary statistics such as the site-frequency spectrum (SFS), we need to augment the state space to keep track of the number of lineages that subtend *i* lineages at a time (Hobolth et al., 2021). Let each state be an *n*-tuple **e** = (*a*_1_, *a*_2_, …, *a*_*n*_) where *a*_*i*_ denotes the number of lineages that subtend *i* lineages in the coalescent tree. Each state is parametrized by *E* = {**e** : Σ_*i*_ *ia*_*i*_ = *n*}, and the only absorbing state is *B* = {(0, 0, …, 1)}, in which the remaining lineage subtends *n* lineages. The transition rates are specified by

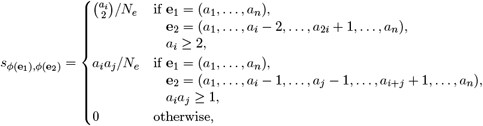

where *ϕ* is an arbitrary ordering, e.g. lexicographic. We refer to this state space as the *block-counting state space*, and to *a*_*i*_ as a *lineage block* (see Appendix B.1.2 for the transition graph).

#### A.5 Rewards

The above transition rates describe the rate of coalescence of any two lineages so that *τ* corresponds to the time to the most recent common ancestor (MRCA) of all lineages. In order to compute more complex summary statistics such as the total branch length and SFS, we need to assign different weights to each state. Let

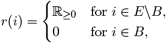

which assigns non-negative values (called rewards) to non-absorbing states, and zero to absorbing states. This gives rise to the reward-transformed absorption time (Hobolth et al., 2024)

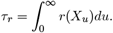

#### A.6 First Moment

The expected reward accumulated over the infinitesimally small interval [*u, u* + *du*], given that the process *X*_*u*_ is in state *k* at time *u*, is *r*(*k*) *du*. By integrating over the expected reward at each time point, we can obtain the reward accumulated over time [*s, t*] given that we start in state *i* and end in state *j*. We have

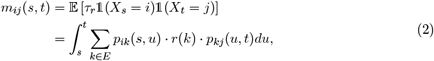

where 1(*X*_*s*_ = *i*) is the indicator function which is 1 when *X*_*s*_ = *i* and 0 otherwise. Here the infinitesimal reward *r*(*k*) *du* is weighted by the probability *p*_*ik*_(*s, u*) of being in state *k* at time *u* given that we started in state *i* at time *s*, and the probability *p*_*kj*_(*u, t*) of being in state *j* at time *t* given that we were in state *k* at time *u*.

We can rewrite equation (2) in matrix form as

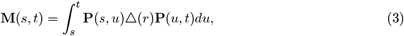

where △(*r*) is a diagonal matrix holding the rewards, and **P**(*s, t*) = {*p*_*ij*_(*s, t*)} represents the transition matrix.

The first moment with respect to the initial state vector ***α*** can then be computed as

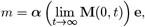

where **e** = (1, 1, …, 1) is a vector of ones, which has the effect of summing the accumulated rewards over all end states.

#### A.7 Time-Inhomogeneity

Assuming that the rate matrix **S** is constant over time, we can compute the transition matrix **P**(*s, t*) simply by taking the matrix exponential:

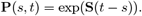

However, determining **P**(*s, t*) for time-inhomogeneous processes where **S** depends on time *u*, requires evaluating a product integral, i.e.,

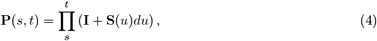

where **I** is the identity matrix (Albrecher and Bladt, 2018).

#### A.8 Van Loan’s Method

The integral described in equation (3) representing the expected reward can be computed using Van Loan’s method (Van Loan, 1978; Hobolth and Jensen, 2011).

**Theorem** (Van Loan’s Method for First Order Moments). *Let*

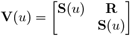

*be a block matrix containing the rate matrix* **S** *and the reward matrix* **R**. *By computing*

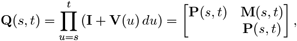

*the accumulated reward* **M**(*s, t*) *over time interval* [*s, t*] *can be obtained from the upper right block of* **Q**(*s, t*).

**Proof Sketch:** To show this, we note that the structure of matrix multiplication and addition and thus product integration implies that we can write

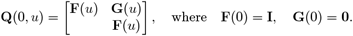

Observing that *d***Q**(0, *u*)*/du* = **V**(*u*)**Q**(0, *u*), we obtain

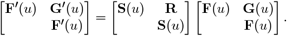

This system of differential equations can be solved to obtain the desired result. We have

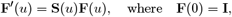

whose solution is (cf. Equation (4))

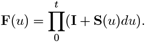

Next, we have

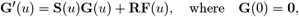

which is solved by (cf. Equation (3))

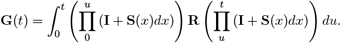

#### A.9 Higher-Order Moments

Van Loan’s method generalizes to higher-order moments (Van Loan, 1978; Hobolth and Jensen, 2011). In order to compute the *k*th (cross-)moment conditioned on the order of rewards, we need to evaluate

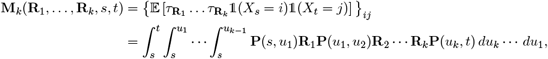

where **R**_*i*_ are the reward matrices.

Let the Van Loan matrix of order *k* be the (*k* + 1) *×* (*k* + 1) block matrix

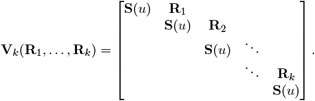

Similar to above, we proceed by taking the product integral of **V**_*k*_, obtaining the (*k* + 1) *×* (*k* + 1) block matrix

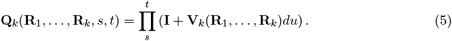

Matrix **M**_*k*_ can then again be obtained from the upper right block of **Q**_*k*_. From this the moment can be computed as

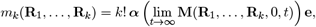

where ***α*** is the initial state vector, and **e** a vector of ones.

The moment *m*_*k*_(**R**_1_, …, **R**_*k*_) is conditioned on the order in which the rewards are accumulated. To obtain the unconditioned moment, we average over all permutations of the reward matrices (cf. Theorem 8.1.5 in Bladt and Nielsen (2017)). Theorem 8.1.5

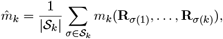

where *S*_*k*_ is the set of permutations of {**R**_1_, …, **R**_*k*_}.

#### A.10 Discretization

In general, product integral (5) is difficult to evaluate precisely for time-inhomogeneous processes. We thus propose a discretization scheme in which the demography is discretized into *l* epochs within which **S** is constant. We can then obtain exact solutions under this discretized demography.

Let **S** = {**S**_*i*_}_*i*_, *i ∈* {0, …, *l ™* 1} be the rate matrices in each epoch:

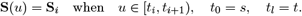

We set up the model such that sufficiently many epochs are generated so as to ensure we have reached almost sure absorption by time *t*_*l*_.

The *k*th moment under a piecewise-constant demography can be obtained by

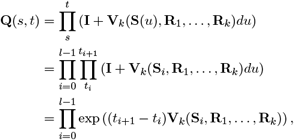

where the last equality follows from the definition of the matrix exponential.

Similarly, to obtain the cumulative distribution function (CDF) for the tree height, we can multiply the transition matrices for each epoch. Let *u ∈* [*t*_*k™*1_, *t*_*k*_) be in the *k*th epoch. The cumulative probability of absorption by time *u* is given by discretizing product integral (4):

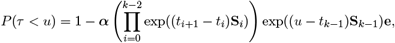

from which we can also obtain the time to almost sure absorption *t*_*abs*_.

### B Transition Rates

#### B.1 Standard Coalescent

The transition rates for the standard coalescent are provided in Section A.4.

##### B.1.1 Lineage-Counting State Space

The state space size is equal to the number of initial lineages *n*, i.e., |*E*| = *n*.

**Figure B4:**
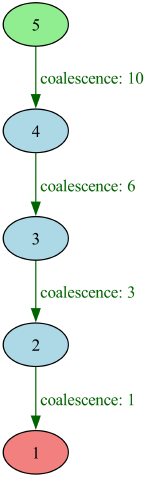
Transition graph for the standard coalescent with five lineages under the lineage-counting state space. Each node represents a state, and each edge a transition. Each state is described by the number of lineages present at the time. The population size *N*_*e*_ is set to 1. Initial states are marked green, and absorbing states red.

##### B.1.2 Block-Counting State Space

The state space size is equal to the number of unordered positive integer partitions of *n*, where *n* is the number of initial lineages. We have

**Figure B5:**
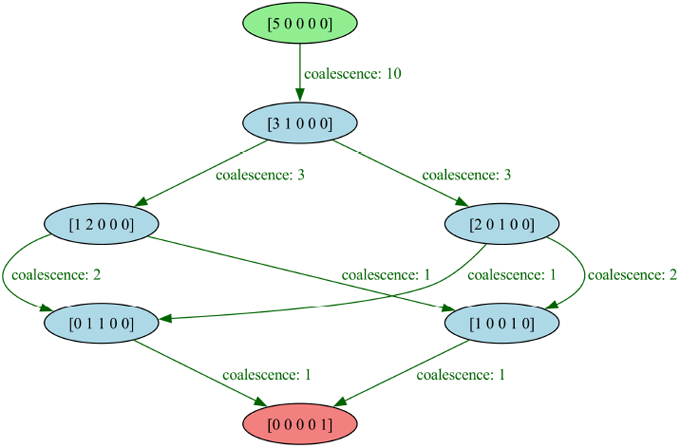
Transition graph for the standard coalescent with five lineages under the block-counting state space. Each state is described by a 5-tuple (*a*_1_, …, *a*_5_) which respresents the number of lineages in each lineage block *a*_*i*_ (cf. Section A.4). The population size *N*_*e*_ is set to 1.

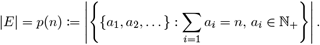

#### B.2 Migration

In this model, we have a finite number of demes, each with its own population size, and which are connected by migration. Migration events are not allowed to occur concurrently with coalescence events, and only one lineage can migrate at a time. The coalescent rates are the same as for the standard coalescent but mergers are only allowed to take place among lineages in the same deme. There exists one absorbing state for each deme, in which the final coalescence event can take place.

##### B.2.1 Lineage-Counting State Space

Let each state be parametrized by **e** = (*d*_1_, …, *d*_*m*_), where *d*_*i*_ is the number of lineages in deme *i*, and *m* is the number of demes. Migration rates are given by

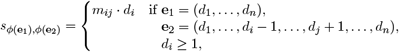

where *m*_*ij*_ is the migration rate from deme *i* to deme *j* backwards in time.

The state space size is given by

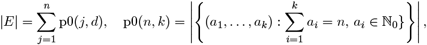

where *n* is the number of initial lineages, *d* the number of demes, and p0 the number of ordered non-negative integer partitions of *n* into *k* parts. The partitions are ordered since demes are not exchangeable, and non-negative since the number of lineages in a deme can be zero.

**Figure B6:**
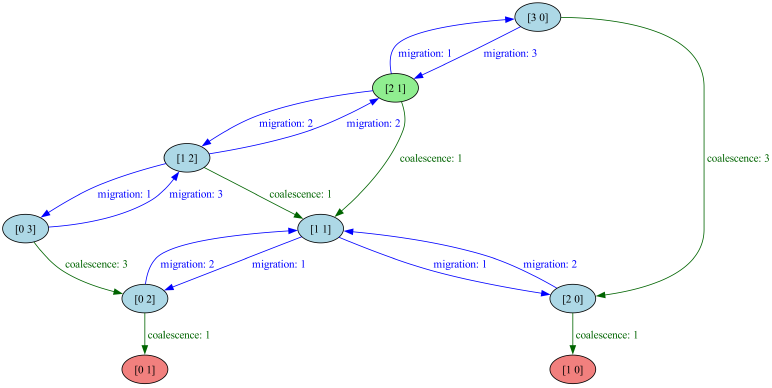
Transition graph for migration in a two-deme model with three lineages under the lineage-counting state space. States are parametrized by 2-tuples respresenting the number of lineages present in each deme. All migration rates and population sizes are set to 1.

##### B.2.2 Block-Counting State Space

Let each state be parametrized by *e* = (**d**_1_, …, **d**_*m*_), where **d**_*i*_ = (*a*_*i*1_, …, *a*_*in*_) are the lineage block counts in deme *i*, and *n* is the initial number of lineages. We require 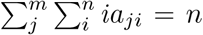, and the migration rates are given by

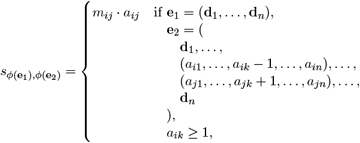

where *m*_*ij*_ is the migration rate from deme *i* to deme *j* backwards in time.

The state space size is given by

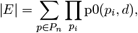

where *n* is the number of initial lineages, *d* the number of demes, *P*_*n*_ the set of unordered positive integer partitions of *n* (cf. Section B.1.2), and p0 the number of ordered non-negative integer partitions of *n* into *k* parts (cf. Section B.1.2).

**Figure B7:**
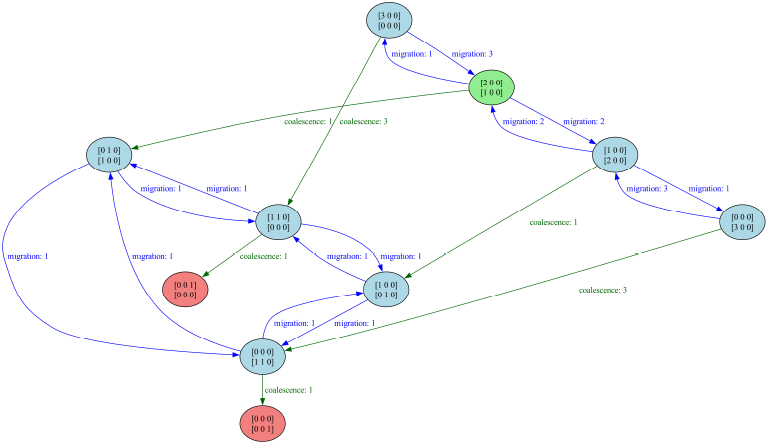
Transition graph for migration in a two-deme model with three lineages under the block-counting state space. States are described by 2-dimensional arrays where the first dimension represents the deme and the second the lineage block counts. All migration rates and population sizes are set to 1. In this setup, the state space is equivalent to the lineage-counting state space as there is only one possible coalescent topology for *n* = 3 lineages.

#### B.3 Beta Coalescent

The beta coalescent is a multiple-merger coalescent where the number of lineages participating in a coalescence event is drawn from a beta distribution parametrized by *α*. Here we scale time by *N*_*e*_ as in the standard coalescent instead of 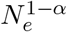 as described in Schweinsberg (2003). The state space is equivalent to that of the standard coalescent, but there are considerably more transitions between states.

##### B.3.1 Lineage-Counting State Space

Let *E* = {1, 2, …, *n*}, where each state describes the number of lineages present, and *n* is the initial number of lineages. Let *α ∈* (1, 2). Then the probability of *k* out of *b* lineages coalescing is given by

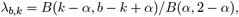

and the transition rates are given by

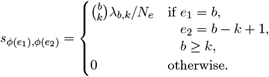

**Figure B8:**
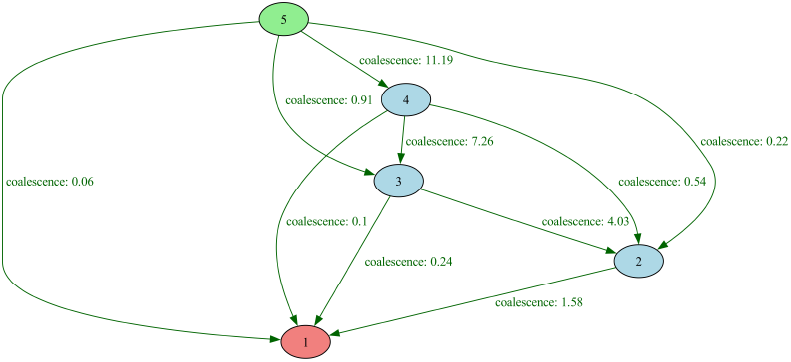
Transition graph for the beta coalescent with five lineages under the lineage-counting state space. The population size *N*_*e*_ is set to 1. Unlike for the standard coalescent, all non-absorbing states are connected to each other.

#### B.3.2 Block-Counting State Space

Let each state be an *n*-tuple **e** = (*a*_1_, *a*_2_, …, *a*_*n*_) such that Σ_*i*_ *ia*_*i*_ = *n* where *a*_*i*_ denotes the number of lineages that subtend *i* lineages in the coalescent tree. Let *k*_*i*_ be the number of lineages in block *i* that coalesce.

The transition rates are given by

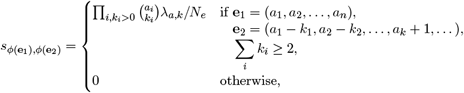

where 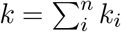 and 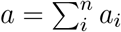.

**Figure B9:**
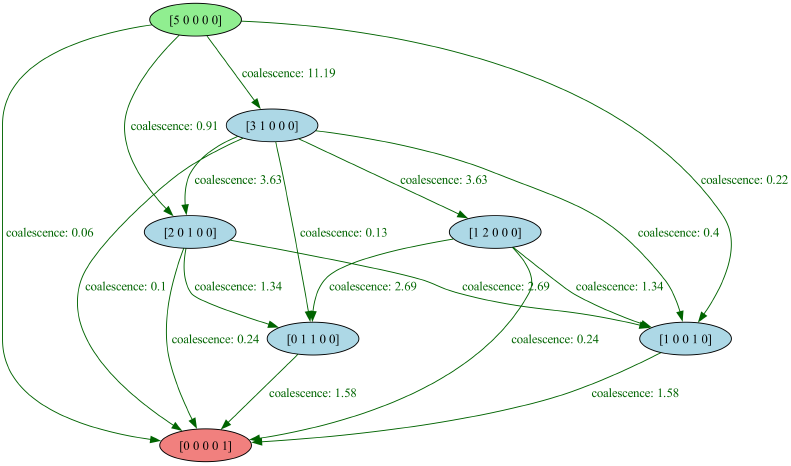
Transition graph for the beta coalescent with five lineages under the block-counting state space. Each state is described by a 5-tuple which respresent the number of lineages in each block. The population size *N*_*e*_ is set to 1.

#### B.4 Dirac Coalescent

The Dirac coalescent is a multiple-merger coalescent where, in addition to binary coalescence events, we allow for multiple-mergers where a fraction *ψ* of the total lineages coalesces with rate *c* (Eldon and Wakeley, 2006; Birkner et al., 2013a). Here we scale time by *N*_*e*_ as in the standard coalescent instead of 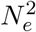 as described in Eldon and Wakeley (2006). The state space is equivalent to that of the standard coalescent, but there are considerably more transitions between states.

**Figure B10:**
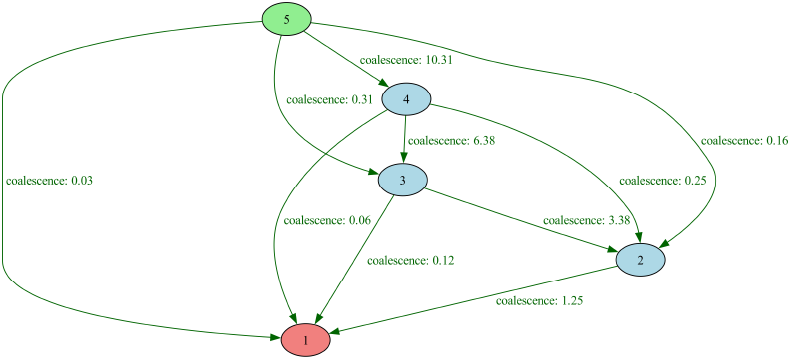
Transition graph for the Dirac coalescent with five lineages under the lineage-counting state space. *N*_*e*_ = 1, *c* = 1, and *ψ* = 0.5.

##### B.4.1 Lineage-Counting State Space

Let *E* = {1, 2, …, *n*}, where each state describes the number of lineages present, and *n* is the initial number of lineages. The transition rates are given by

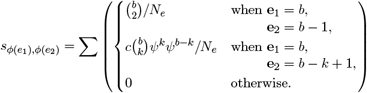

##### B.4.2 Block-Counting State Space

Let each state be an *n*-tuple **e** = (*a*_1_, *a*_2_, …, *a*_*n*_) such that _*i*_ *ia*_*i*_ = *n* where *a*_*i*_ denotes the number of lineages that subtend *i* lineages in the coalescent tree. Let *k*_*i*_ be the number of lineages in block *i* that merge. The transition rates are given by

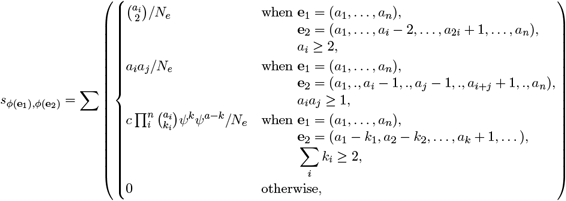

where 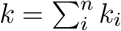 and 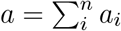.

**Figure B11:**
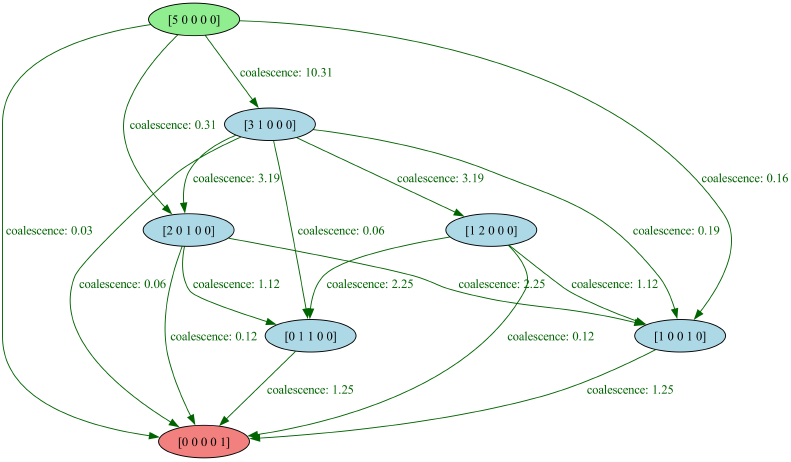
Transition graph for the Dirac coalescent with five lineages under the block-counting state space. Each state is described by a 5-tuple which respresent the number of lineages in each block. We have *N*_*e*_ = 1, *c* = 1, and *ψ* = 0.5.

#### B.5 Recombination

Here we consider recombination between two loci under the standard coalescent and lineage-counting state space. Extensions are possible but the state space grows extremely rapidly with the number of loci, and under the block-counting state space. Allowing for multiple demes is straightforward as migration and recombination events occur independently of each other.

##### B.5.1 Lineage-Counting State Space

Let each state be parametrized by **e** = (*l*_12_, *l*_1_, *l*_2_), where *l*_12_ is the number of linked lineages, i.e. shared by both loci, and *l*_*i*_, *i ∈* {1, 2} are the number of unlinked lineages at locus *i*, i.e. not shared with the other locus. A recombination event reduces the number of linked lineages by one, and increases the number of unlinked lineages by one:

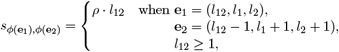

where *ρ* is the recombination rate.

Four types of coalescence events can occur. In a *linked coalescence* event, two linked lineages coalesce:

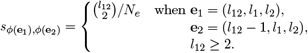

Similarly, in an *unlinked coalescence* event, two unlinked lineages from the same locus coalesce:

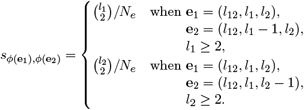

We may also have *mixed coalescence* events where an unlinked lineage from one locus coalesces with a linked lineage:

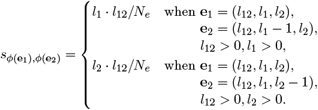

Finally, we may also have *locus coalescence* events where two unlinked lineages from different loci coalesce:

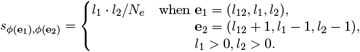

When considering more than two lineages, mixed and unlinked coalescence events can both occur for the same transition configuration (*e*_1_, *e*_2_).

The state space size equals the maximum sum of products of successive pairs in a permutation of order *n* + 1 for *n >* 1. We have

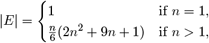

where *n* is the number of initial lineages.

**Figure B12:**
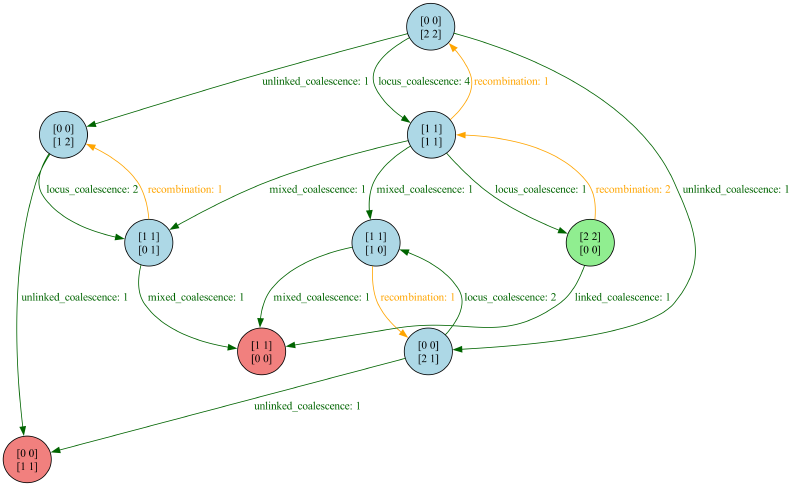
Transition rates for 2-locus recombination with two lineages under the standard coalescent and lineage-counting state space. Each state is parametrized by two 2-tuples which respresent the number of linked lineages per locus (top part) and the number of unlinked lineages per locus (bottom part). In the starting state, both lineages are linked (cf. green node). We have 2 obsorbing states, one in which both remaining lineages are linked and one in which they are unlinked.

### C Code Examples

#### C.1 Complex Code Example

In this code example, we define a Coalescent distribution for a two-deme model with 8 lineages, using the beta coalescent model (Schweinsberg, 2003). We specify a 3-epoch demography, involving a bottleneck in one of the demes, and a single migration rate change. Note that the keys and values in the demography dictionaries represent the time of change and the corresponding population size or migration rate, respectively.

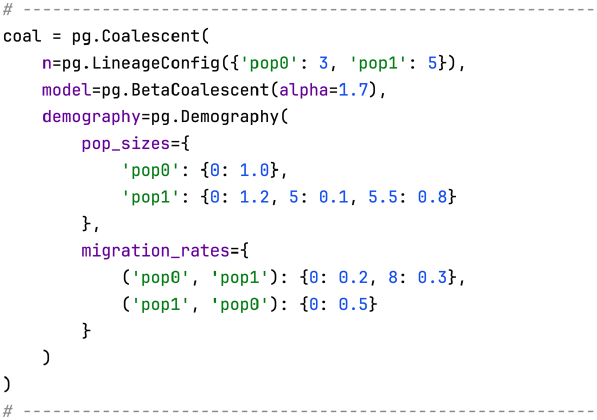

Below, we proceed to plot several quantities, specifically the demography, tree height distribution, expected SFS, and 2-SFS correlation matrix. Note that the maximum time displayed is adjusted to the 99th percentile of the tree height distribution. The output of the following code snippet is shown in Figure C13 (cf. plot_manuscript_complex_example.py).

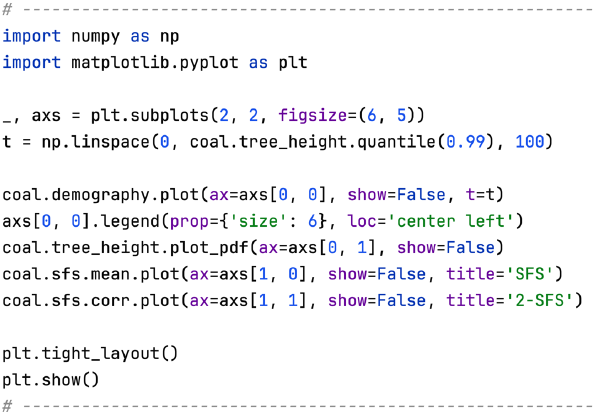

**Figure C13:**
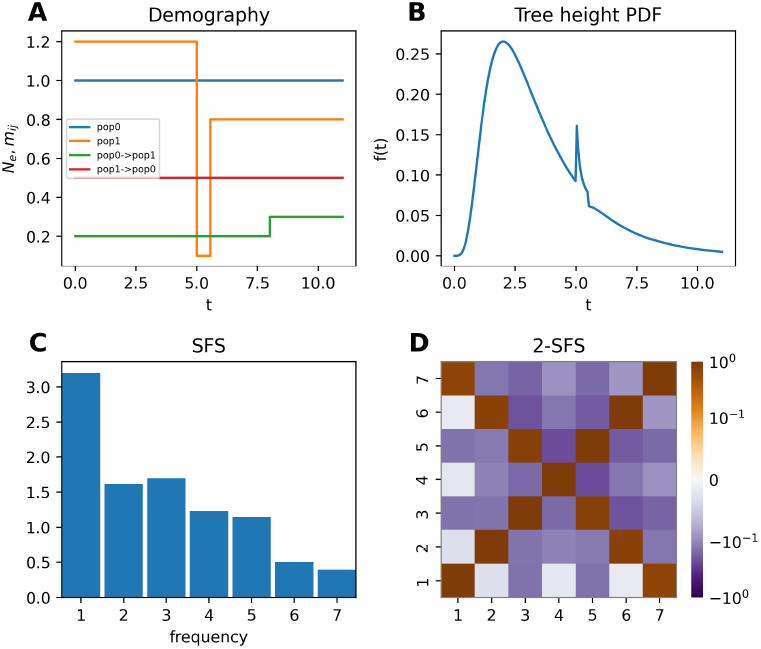
Output from code snippet in Section C.1. **A**: population sizes and migration rates over time. **B**: tree height distribution. We observe a temporary peak in the coalescent rate during the bottleneck. **C**: expected SFS. There appears to be a relative deficit of doubletons under this demographic scenario. **D**: 2-SFS correlation matrix representing the branch length correlation of branches subtending i and j lineages in the coalescent tree.

### D State Space Size

State space sizes for different numbers of lineages and demes are shown in Figures D14 and D15. Note that the size of the resulting Van Loan matrix to be exponentiated is *m*(*k* + 1) where *m* is the number of states and *k* the order of the moment of interest.

**Figure D14:**
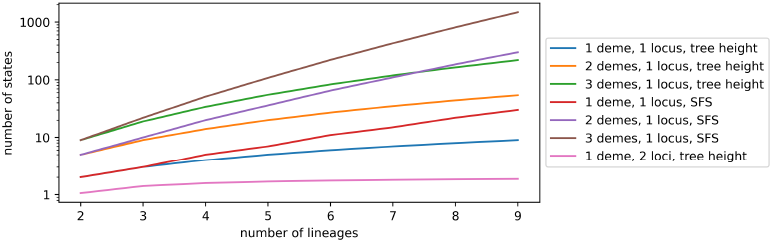
State space size for different numbers of lineages (x-axis), with varying numbers of demes and loci as shown in the legend. For summary statistics based on the tree height the lineage-counting state space is sufficient, while for SFS-based statistics the block-counting state space is required.

**Figure D15:**
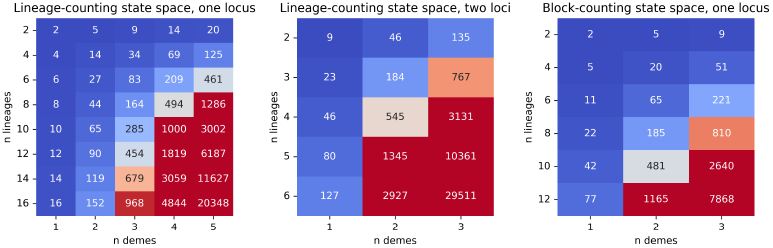
State space size for different numbers of lineages and demes. We consider the lineage-counting state space with one (left) and two loci (middle), as well as the block-counting state space with one locus (right). The state space size grows exponentially with the number of lineages and demes, which can heavily impact computation time (Section E).

### E Computation Time

Computation times for different numbers of lineages and demes are shown in Figures E16 and E17. Keeping the state space small is crucial for maintaining reasonable computation times (e.g., under a few minutes for a few thousand states). In general, state spaces with a few hundred states can be computed quickly and are well-suited for gradient-based optimization. Total runtimes depend on several factors: the complexity of the demographic model, i.e., the number of lineages, loci, and demes; the number of epochs; the order of the moment of interest; the type of summary statistic; the type of state space; and, for parameter estimation, the number of parameters to be estimated. The most limiting factor is the size of the state space, as the runtime complexity of matrix exponentiation scales roughly cubically with the number of states. For summary statistics based on the tree height (including the CDF, PDF, and quantile function) or total branch length, the lineage-counting state space is sufficient, which only grows linearly with the number of lineages (cf. Section D). For statistics based on the SFS, the block-counting state space is required, which grows exponentially both with the number of lineages and demes. In addition, *n ™* 1 independent matrix exponentiations are needed for the SFS, where *n* is the number of initially present lineages.

If not the entire SFS is required, it is thus advisable to compute moments in a more targeted way (i.e., by using Coalescent.moment rather than Coalescent.sfs.mean). All branch length–based summary statistics require exponentiating the Van Loan matrix, which has size *m*(*k* + 1), where *m* is the number of states and *k* the order of the moment (cf. Section A.8). Upon creation and initialization of the Coalescent object, warnings are issued if the state space size exceeds certain thresholds. Alternatively, the user can directly retrieve the respective state space size by accessing [Coalescent.lineage_counting_state_space.k_or_Coalescent.block_counting_state_space.k](https://phasegen.readthedocs.io/en/v1.0.1/modules/state_space.html#phasegen.state_space.BlockCountingStateSpace.k). The computation time is linear in the number of epochs, however, since the order of the Van Loan matrix remains constant; instead *l* Van Loan matrices are multiplied together, where *l* is the number of epochs. By default, matrix exponentiation is performed using double-precision floatingpoint numbers, which provides very accurate results. Some performance improvements can be achieved by using single-precision floating-point numbers, but this may lead to numerical issues in some cases. Finally, when performing parameter estimation, the complexity of the loss landscape increases rapidly with the number of parameters. This also depends on the optimization algorithm; by default, the `L-BFGS-B` algorithm is used, but alternatives may be specified.

In contrast, `dadi`’s runtime remains largely unaffected by the SFS sample size, as it relies on discretizing the same underlying continuous differential equation. However, long integration times and severe population size reductions forward in time can increase computation times, whereas `PhaseGen` is largely unaffected by these factors. Since both `dadi` and `PhaseGen` use similar numerical optimization procedures, their performance scales similarly with the number of parameters being estimated. Nevertheless, as model complexity increases, `PhaseGen` becomes significantly slower than `dadi`.

**Figure E16:**
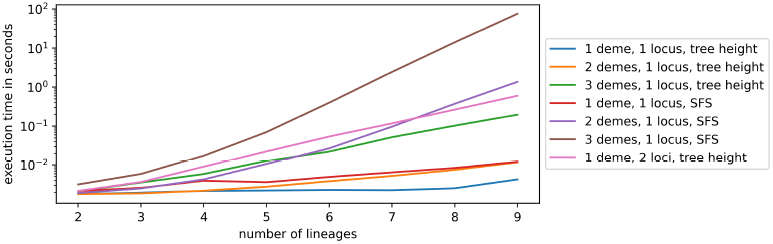
Total computation time (in seconds) for calculating expected tree height and SFS across varying numbers of lineages (x-axis), with different demes and loci as indicated in the legend. Computations were performed on a single core of a MacBook Pro M2 using the `SciPyExpmBackend` for matrix exponentiation. Larger state spaces may be computed more efficiently using alternative backends (https://phasegen.readthedocs.io/en/v1.0.1/modules/expm.html). The computation time of matrix exponentiation is rougly cubic in the number of states, depending on the structure of the matrix and algorithms used.

**Figure E17:**
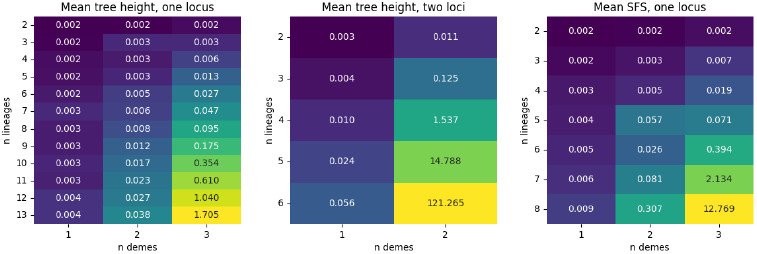
Total computation time (in seconds) for different numbers of lineages, demes and loci using the `SciPyExpmBackend` for matrix exponentiation. We consider the lineage-counting state space (left and middle), and the block-counting state space (right). Note that the computation of the SFS requires *n ™* 1 independent matrix exponentiations, where *n* is the number of lineages initially present.

### F Comparison against msprime results

To validate our implementation, we compare the results from `PhaseGen` to those from `msprime`. To this end, we fully reimplement the Coalescent class, obtaining results by averaging over many coalescent trees simulated with `msprime`, rather than relying on phase-type theory. By simulating a sufficiently large number of trees (typically 10^6^), we obtain accurate estimates, which we compare to the results from `PhaseGen` using a strict deviation threshold. In total, we compare 81 different scenarios, including the standard, beta, and Dirac coalescents; 1 or 2 loci; varying numbers of demes and lineages; different recombination and migration rates; changing migration and population sizes; and varying numbers of epochs. We also include extreme cases, such as rates differing by several orders of magnitude and highly dynamic demographic events like strong bottlenecks, to test for potential numerical issues. All results were consistent with expectations, with no significant discrepancies observed, even in more extreme cases. In addition, mechanisms are in place to warn users about potential numerical issues and mis-specified demographies that result in an infinite time to the most recent common ancestor (TMRCA). Tested statistics include the tree height CDF and PDF; the first four moments of tree height, total branch length, and the SFS (folded and unfolded); the 2-SFS; centered moments such as the variance and 2-SFS covariance; and moments marginalized over demes and loci. The configuration files specifying the scenarios, along with the tolerance thresholds for the tested statistics, can be found in the `resources/configs` directory of the `PhaseGen` repository. In addition, all unit tests are located in the `testing` directory.

